# Maternal exercise rescues fetal akinesia-impaired joint and bone development

**DOI:** 10.1101/2025.06.17.660083

**Authors:** Christopher J. Panebianco, Yuming Huang, Nidal Khatib, Devin C. Gottlieb, Maha Essaidi, Saima Ahmed, Nathaniel A. Dyment, Rebecca A. Simmons, Joel D. Boerckel, Niamh C. Nowlan

**Author notes:** Corresponding authors: Joel D. Boerckel, University of Pennsylvania, Department of Orthopaedic Research, Department of Bioengineering, 376A Stemmler Hall, 3450 Hamilton Walk, Philadelphia, PA 19104-6081 (Email), Niamh C. Nowlan, University College Dublin, UCD School of Mechanical and Materials Engineering, Belfied, Dublin 4, Ireland (Email).

## Abstract

Fetal movements exert mechanical forces that shape the developing skeleton. Conditions that impair fetal movements can cause skeletal defects, but interventions are limited. Here, we show that maternal wheel running exercise regulates fetal skeletal development in mice. In wild-type fetuses, maternal exercise stimulated joint and bone morphogenesis. We reasoned that these effects occurred through either indirect maternofetal communication or direct mechanical stimulation of the fetus. Maternal exercise did not alter placental measures of nutrient transport. However, in the Splotch-delayed (Sp^d^) mouse model of fetal akinesia, which features intact maternofetal communication but lacks fetal movements, maternal exercise substantially rescued fetal akinesia-impaired joint and bone development and prevented disuse-induced resorption of the deltoid tuberosity. Further, direct mechanical stimulation of Sp^d^ limbs explanted from systemic factors similarly stimulated joint morphogenesis. Together, these findings identify maternal exercise as a regulator of fetal skeletal development, providing a platform for studying skeletal developmental mechanobiology and suggesting potential therapeutic applications for fetuses with impaired movement.

## 1.0 Introduction

Spontaneous fetal movements exert mechanical forces that shape the developing skeleton. The first fetal movements begin with the initiation of muscle formation and coincide with joint and bone development.^1,2^ Reduced fetal movements (*i.e.,* fetal hypokinesia) can cause congenital defects in skeletal development, including hip dysplasia, congenital scoliosis, joint contractures, and delayed bone formation.^3,4^ Etiologically, fetal hypokinesia-associated syndromes exist on a broad phenotypic spectrum from mild to perinatal lethal, with fetal akinesia (*i.e.,* no movements) being the most severe.^5^ Fetal hypokinesia can be associated with intrauterine environmental conditions, such as breech fetal position^6,7^ and insufficient amniotic fluid (*i.e.,* oligohydramnios),^8^ genetic mutations that affect neuromuscular development (*e.g.,* genetic forms of arthrogryposis),^9^ or non-genetic conditions with abnormal movements *in utero* (*e.g.,* amyoplasia).^10,11^ While fetal hypokinesia-causing conditions can sometimes be detected by prenatal imaging and/or genetic screening, treatment of skeletal abnormalities is currently limited to postnatal interventions, including surgeries, bracing, and physical therapy.^12–17^ This treatment gap stems both from limited mechanistic understanding of how fetal movements direct skeletal development and limited means for prenatal intervention. Here, using mouse models, we demonstrate that maternal exercise during fetal gestation can alter fetal joint and bone morphogenesis and can partially rescue both joint and bone formation in a preclinical model of fetal akinesia.

Animal models of limited fetal movement recapitulate the skeletal defects observed in fetal akinesia patients. Early experiments used transplanted embryonic chick limb buds, which lack innervation, to infer that disrupted muscle development causes defects in joint cavitation and shape and bone formation.^18,19^ Subsequent studies used pharmacological agents and motor neuron resection to paralyze chick embryos, and observed similar defects in both joint and long bone development.^20–28^ Mouse models of fetal akinesia have further enabled mechanistic insights into how muscle contractions guide mammalian skeletal development. The muscular dysgenesis (*mdg*) and Splotch-delayed (Sp^d^) mouse models are products of spontaneous mutations. *Mdg* mice harbor a mutation in the DHPɑR1 receptor that prevents excitation-contraction coupling^29–31^ and Sp^d^ mice feature a point mutation at the Pax3 gene that abrogates muscle progenitor cell migration and limits skeletal muscle development.^32–34^ Additional models have been genetically engineered to disrupt fetal muscle development, including the *Myf5*^-/-^:*MyoD^-/-^* amyogenic mouse, which lacks striated muscle due to deletion of the myogenic regulatory factors *Myf5* and *MyoD*.^35^ Skeletal development studies using these models consistently show that *in utero* muscle function is required for proper fetal joint morphogenesis and bone formation.^36–43^ Additionally, fetal muscle contraction regulates the growth and maturation of bony eminences at tendon insertion sites, such as the deltoid tuberosity of the humerus, which initiates via a cartilaginous rudiment that resorbs in the absence of fetal movements.^44^ The coherence of these findings across human patients and animal models implicates the loss of muscle-generated mechanical forces in fetal akinesia-associated defects in skeletal development.

*Ex vivo* bioreactor cultures complement *in vivo* animal models of fetal akinesia by allowing the application of direct mechanical stimulation during fetal skeletal development.^45–48^ Previously, using bioreactor culture of embryonic chick limb explants, we showed that the amplitude, frequency, and duration of joint flexion regulate joint morphogenesis *ex vivo*.^49,50^ We further used mouse limb explant culture to demonstrate the roles of specific molecular mechanosensors and mechanotransducers in the mechanoregulation of skeletal morphogenesis. Using pharmacologic inhibition and conditional genetic ablation approaches, we showed that the mechanosensitive ion channel, Transient receptor potential cation channel subfamily V member 4 (TRPV4), mediates joint morphogenesis in response to applied limb movement,^51,52^ and the mechanoresponsive transcriptional regulators, Yes-associated protein (YAP) and Transcriptional co-activator with PDZ-binding motif (TAZ), mediate load-induced fetal bone formation.^53^ Developmentally, we found that YAP/TAZ mechanosignaling in vessel-associated osteoblast precursor cells coordinates blood vessel invasion and growth plate remodeling at the chondro-osseous junction during fetal bone development.^53^ These findings suggest that both cartilaginous joint morphogenesis and endochondral ossification are mechanically regulated. In the context of fetal akinesia, we found that *ex utero* dynamic mechanical stimulation rescues collagen extracellular matrix deposition and organization in Sp^d^ mice.^54^ These bioreactor studies have enabled interrogation of the molecular basis by which absent and restored mechanical signals regulate skeletal morphogenesis, but are limited by extraction from the fetal vascular system and from systemic factors provided by the gestational environment. Thus, explants can only be cultured for a limited duration in an environment that cannot perfectly mimic *in utero* conditions.

In addition to active movements exerted by the fetus, passive movements imparted on the fetal skeleton by maternal activity may also guide development. Motivated by the observation that Sp^d^ and *mdg* mice exhibit greater defects in the forelimb than hindlimb,^55–57^ we reasoned that passive mechanical stimuli from maternal movements may produce distinct mechanical environments in forelimbs and hindlimbs. By applying displacement boundary conditions, such as could be brought about by maternal activity, to finite element models of the forelimb and hindlimb, we observed greater stresses and strains in the developing femur compared to the humerus.^58^ Based on this finding, we hypothesized that passive fetal movements induced by maternal exercise could provide formative mechanical signals to the developing fetus. Here, we test this hypothesis through application of maternal exercise in wild-type and Sp^d^ mice, with a focus on morphogenesis of the humerus.

Current guidelines recommend maternal exercise for healthy pregnancies. Regular moderate-intensity exercise during pregnancy reduces maternal risk of gestational disorders (*e.g.,* gestational diabetes, gestational hypertension),^59–61^ enhances birth outcomes (*e.g.,* reduced preterm birth rate, increased vaginal birth rate),^62–64^ and improves offspring health (*e.g.,* cardiac function, cerebral maturation).^65,66^ Though clinical evidence suggests the multifaceted benefits of maternal exercise, we do not fully understand how maternal exercise affects fetal development. Prior studies suggest that maternal exercise may increase nutrient transport to the developing fetus via the placenta and/or upregulate specific humoral signaling molecules that are transferred to the developing fetus (*e.g.,* insulin-like growth factor 1 (IGF-1)).^67–69^ However, experimental evidence is limited and continued research is required.

Here, we show that maternal exercise regulates fetal skeletal development in mice. We find that maternal exercise stimulates joint morphogenesis, deltoid tuberosity formation, and bone growth in wild-type fetuses and largely rescues fetal akinesia-impaired skeletal abnormalities in Sp^d^ mice. Our data suggest that these benefits are mediated by passive movements imparted on the developing skeleton by maternal exercise, but do not rule out contributions from altered maternofetal signaling. These findings identify maternal exercise as a regulator of fetal skeletal development, implicate maternal exercise as a putative platform for studying skeletal developmental mechanobiology *in vivo*, and motivate further investigation of maternal exercise as a potential *in utero* intervention for fetal hypokinesia-associated skeletal syndromes.

## 2.0 Results

### 2.1 Effects of maternal exercise on fetal joint and bone morphogenesis

We first assessed the effects of maternal exercise on embryonic day 17.5 (E17.5) mouse elbow joint morphogenesis using optical projection tomography (OPT) (**Figure 1A-C**). Pregnant female dams were exercised daily starting at E13.5, which coincides with the onset of fetal limb movements and the early stages of endochondral ossification.^1,2^ Sham controls had the same wheel access; however, their wheels were locked to prevent running (**Figures 1A & S1A**). Maternal exercise increased humeral medial condyle width, height, and depth (**Figure 1D-F**), lateral condyle width, height, and depth (**Figure 1G-I**).

**Figure 1.**
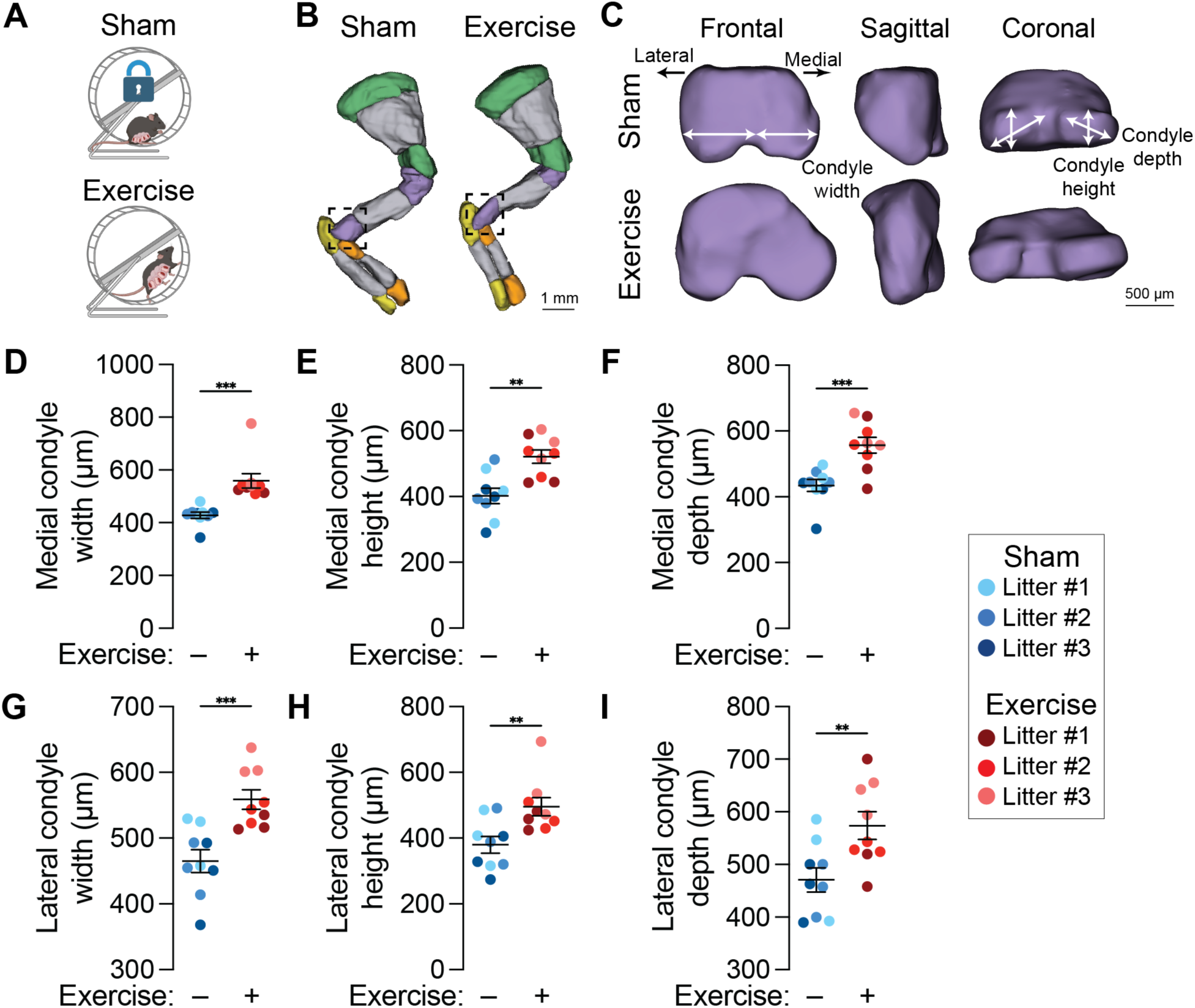
Effects of maternal exercise on fetal joint morphogenesis. **(A)** Supervised wheel running exercise (Exercise) vs. locked wheel controls (Sham). Female mice were acclimated with *ad libitum* running wheel access for two weeks prior to timed pregnancy. Pregnant dams then had no wheel access until embryonic day 13.5 (E13.5). At E13.5, pregnant dams were assigned to Sham or Exercise groups. Exercised mice performed four bouts of 15 minutes of supervised wheel running daily until E17.5. **(B)** Optical projection tomography (OPT) reconstructions of alizarin red and alcian blue-stained forelimbs at E17.5. Cartilaginous regions of the scapula, humerus, radius, and ulna are marked in green, purple, orange, and yellow, respectively. Ossified regions are marked in gray. Scale bar = 1 mm. **(C)** Zoomed projections of the cartilaginous distal humerus. The width, depth, and height of the lateral condyle and medial condyle are marked in the frontal, sagittal, and coronal planes. Scale bar = 500 µm. **(D)** Medial condyle width, **(E)** medial condyle height, **(F)** medial condyle depth, **(G)** lateral condyle width, **(H)** lateral condyle height, and **(I)** lateral condyle depth quantifications. Fetuses from the same litter are marked by the same color datapoint. * = *p*<0.05 using a Student’s t-test.

To test the effects of maternal exercise on bone morphogenesis, we performed OPT and microcomputed tomography (µCT) imaging (**Figure 2A&G**). Maternal exercise significantly increased humerus rudiment length and mineralized length of the bony primary ossification center, but did not affect the mineralization ratio (*i.e.*, the ratio of mineralized length to rudiment length) (**Figure 2B-D**). This indicates that maternal exercise proportionally stimulated rudiment growth and mineralization. The development of the deltoid tuberosity has been shown to be particularly sensitive to absent muscle stimulation.^44^ Maternal exercise significantly increased deltoid tuberosity volume (**Figure 2E**), even after normalization by the rudiment length (**Figure 2F**). Using µCT, we confirmed that maternal exercise increased humeral bone volume and bone length (**Figure 2G-I)**, and further showed that maternal exercise significantly increased bone tissue mineral density (TMD) (**Figure 2J**).

**Figure 2.**
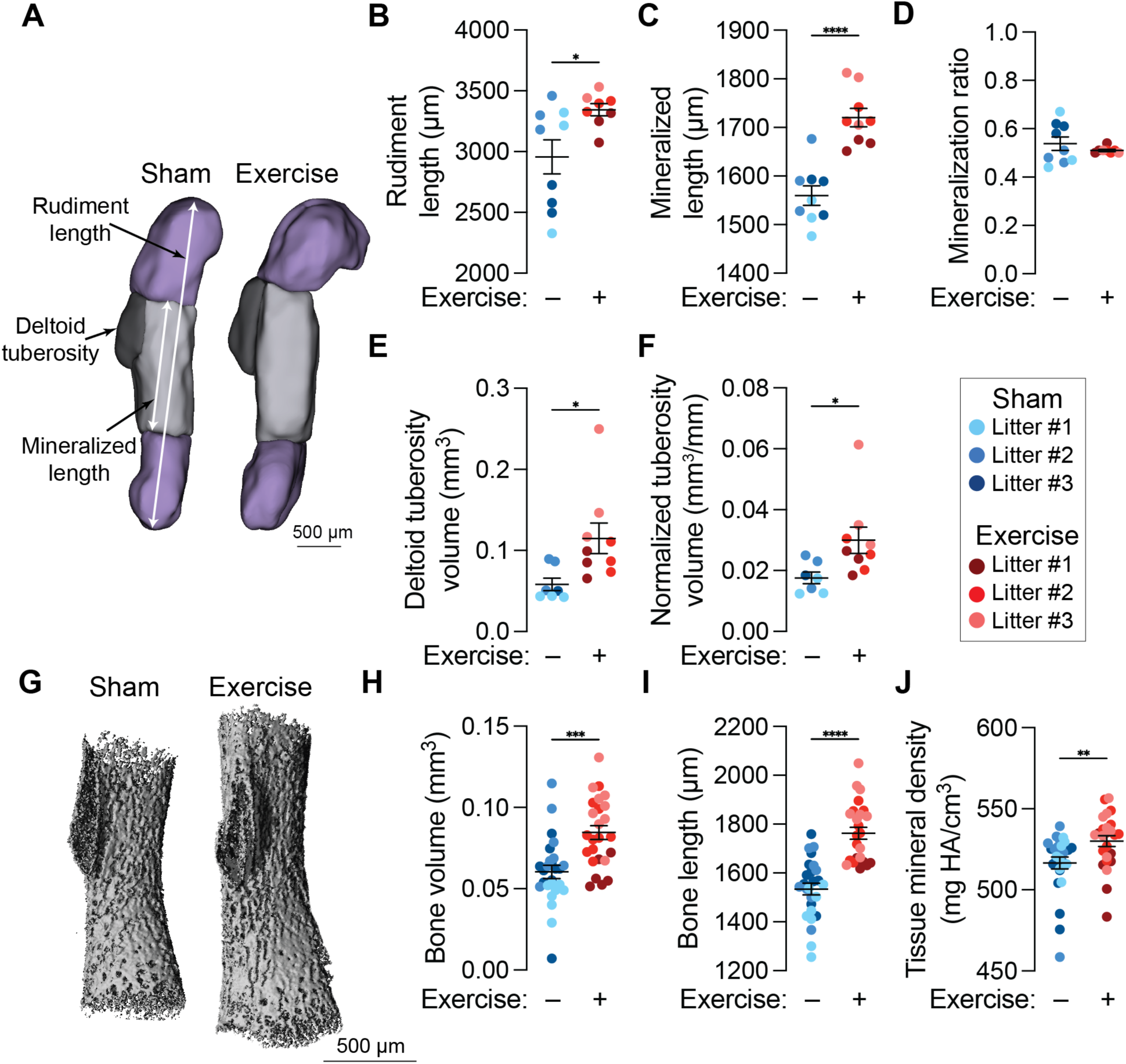
Effects of maternal exercise on fetal bone development. **(A)** Optical projection tomography (OPT) reconstructions of embryonic day 17.5 (E17.5) humeri. The rudiment length and mineralized length are marked with white arrows. The deltoid tuberosity is contoured in black. Scale bar = 500 µm. **(B)** Rudiment length, **(C)** mineralized length, and **(D)** mineralized ratio quantifications (i.e., the ratio of mineralized length to rudiment length). **(E)** Deltoid tuberosity volume and **(F)** rudiment length-normalized deltoid tuberosity volume quantifications. **(G)** Three-dimensional microcomputed tomography (µCT) reconstructions of the E17.5 humeri. Scale bar = 500 µm. **(H)** Bone volume, **(I)** bone length, and **(J)** tissue mineral density (TMD) quantifications. Fetuses from the same litter are marked by the same color datapoint. * = *p*<0.05 using a Student’s t-test.

### 2.2 Effects of maternal exercise on placental transport

Maternal exercise may affect fetal development by affecting the transport of factors or nutrients across the placenta. The fetal weight to placental weight (FW/PW) ratio is an approximate metric of placental transport efficiency, which can be altered by changes in fetal weight, placental weight, or both. A smaller placenta that can efficiently support a larger fetus (*i.e.,* greater FW:PW ratio) suggests greater placental transport efficiency.^70,71^ Maternal exercise significantly increased fetal weight and decreased placental weight, resulting in a significantly increased FW:PW ratio (**Figure 3A-C**). This suggests that maternal exercise may enhance placental nutrient transport to the developing fetus. To further test this hypothesis, we measured the area ratio between the labyrinth zone (LZ) and junctional zone (JZ) of the placenta as a placental metric of transport efficiency. Increased LZ:JZ ratios indicate increased placental transport efficiency.^70,71^ Maternal exercise did not significantly alter the LZ:JZ ratio (**Figure 3D-E**).

**Figure 3.**
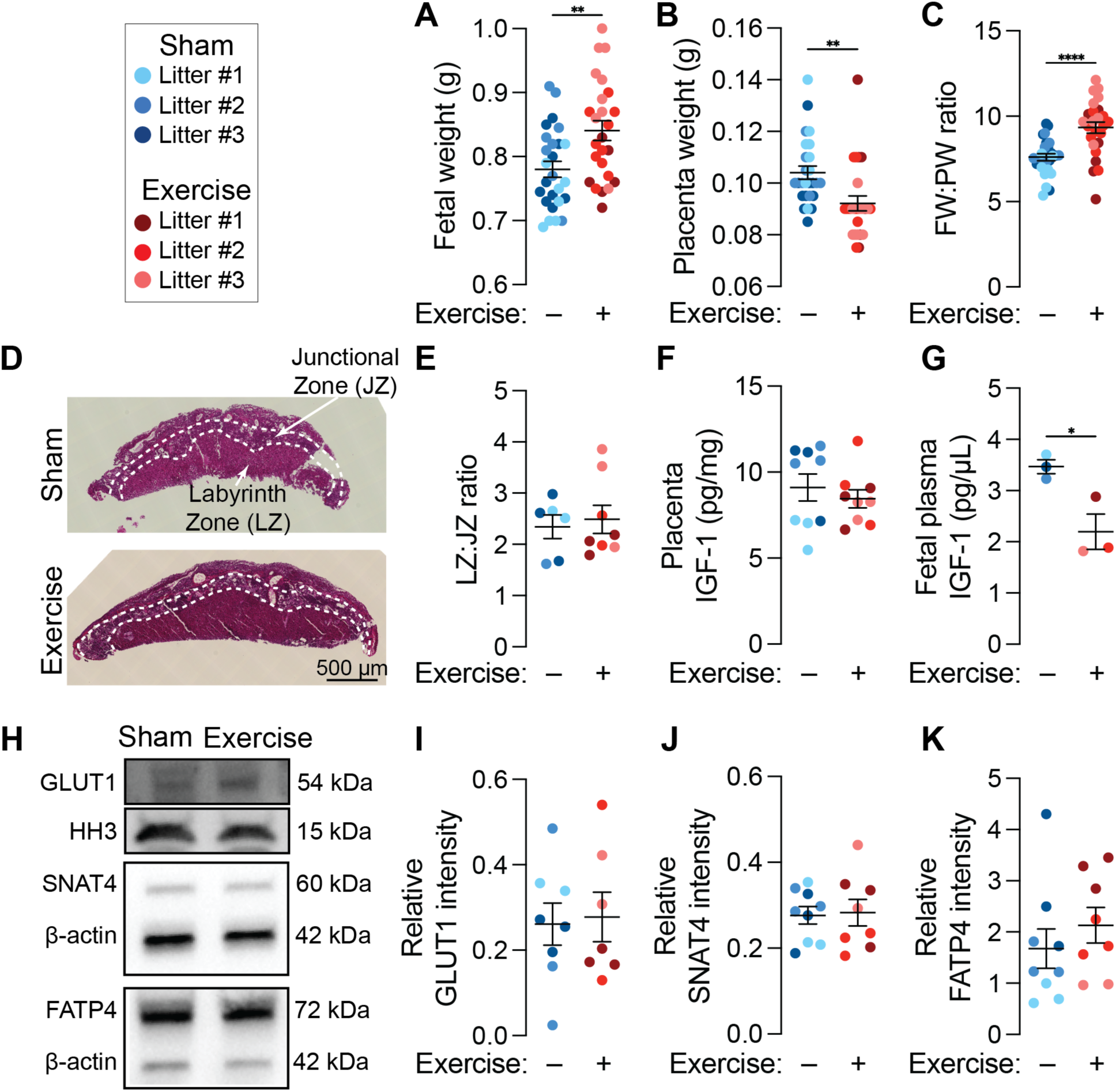
Effects of maternal exercise on placental transport. **(A)** Fetal weight (FW), **(B)** placental weight (PW), and **(C)** FW:PW ratio quantifications. **(D)** Hematoxylin and eosin micrographs of embryonic day 17.5 (E17.5) placentas. Scale bar = 500 µm. **(E)** Labyrinth zone (LZ) area to junctional zone (JZ) area ratio (LZ:JZ) quantifications. **(F)** Placental levels of insulin-like growth factor-1 (IGF-1) and **(G)** fetal plasma levels of IGF-1 quantifications. **(H)** Western blots for glucose transporter 1 (GLUT1, molecular weight (MW) = 54 kDA), sodium-coupled neutral amino acid transporter 4 (SNAT4, MW = 60 kDa), and fatty acid transporter 4 (FATP4, MW = 72 kDa). Histone H3 (MW = 15 kDa) and *β*-actin (MW = 42 kDa) were used as loading controls. **(I)** Relative GLUT1 intensity, **(J)** relative SNAT4 intensity, and **(K)** relative FATP4 intensity quantifications. Fetuses from the same litter are marked by the same color datapoint. * = *p*<0.05 using a Student’s t-test.

Next, we conducted molecular assays of placental transport. Insulin-like growth factor-1 (IGF-1) is the major upstream regulator of placental nutrient transporter expression and IGF-1 levels are strongly correlated with fetal growth outcomes.^72,73^ Maternal exercise did not significantly alter placental levels of IGF-1, but significantly decreased the concentration of IGF-1 in fetal plasma (**Figure 3F-G**). The most abundantly expressed placental nutrient transporters for glucose, amino acids, and fatty acids are glucose transporter 1 (GLUT1), sodium-coupled neutral amino acid transporter 4 (SNAT4), and fatty acid transporter 4 (FATP4), respectively. Consistent with our LZ:JZ and placental IGF-1 findings, maternal exercise did not alter placental levels of GLUT1, SNAT4, or FATP4 (**Figure 3H-K**).

### 2.3 Mechanoregulation of endochondral ossification via maternal exercise

Beyond placental transport, maternal exercise may also stimulate skeletal development through direct mechanical stimulation. Based on our prior studies on the mechanotransductive roles of the transcriptional regulators Yes-associated protein (YAP) and Transcriptional co-activator with PDZ-binding motif (TAZ) in bone development,^53^ we evaluated YAP expression and the abundance of the canonical YAP target gene, Cysteine-rich angiogenic inducer 61 (Cyr61), in the developing bone. YAP is transcriptionally active in the nucleus, and its cytoplasmic-nuclear transport can be induced by a variety of signals, including mechanical cues and morphogen signaling.^74^ Maternal exercise did not significantly alter nuclear YAP intensity in the bone collar (BC) or primary ossification center (POC) (**Figure 4A-C**), but significantly increased Cyr61 abundance in the bone collar (**Figure 4D**). The increase in Cyr61 abundance in the primary ossification center was not statistically significant (*p* = 0.058) (**Figure 4E**).

**Figure 4.**
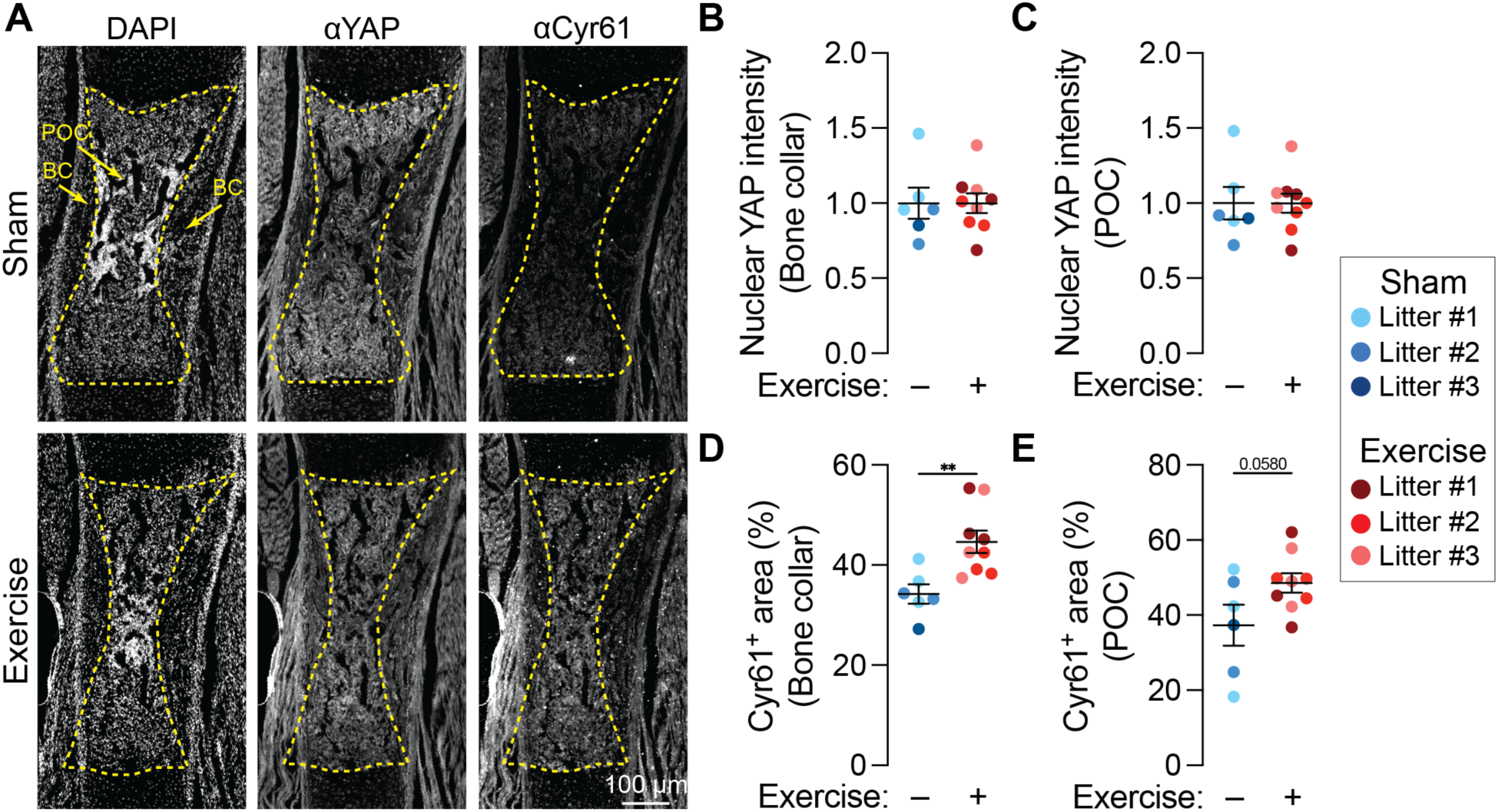
Effects of maternal exercise on YAP signaling. **(A)** Fluorescent staining for nuclei (DAPI), Yes-associated protein (YAP), and Cysteine-rich angiogenic inducer 61 (Cyr61) in embryonic day 17.5 (E17.5) humeri. Scale bar = 100 µm. **(B)** Bone collar (BC) nuclear YAP intensity, **(C)** Primary ossification center (POC) nuclear YAP intensity, **(D)** BC Cyr61-positive area, and **(E)** POC Cyr61-positive area quantifications. Fetuses from the same litter are marked by the same color datapoint. * = *p*<0.05 using a Student’s t-test.

Next, we evaluated the effects of maternal exercise on growth plate morphometry, alkaline phosphatase (ALP) activity, and neovascular morphometry in the developing humerus. These analyses were prioritized based on our findings that osteoprogenitor cell YAP/TAZ signaling regulates fetal bone development by coupling osteoprogenitor mobilization to blood vessel invasion for chondro-osseous junction remodeling and bone formation.^53^ Maternal exercise significantly reduced hypertrophic zone length (**Figure 5A-B**), consistent with increased chondro-osseous junction remodeling.^53^ Maternal exercise did not alter ALP activity, which marks osteogenic cell activity, in either the endochondral primary ossification center or the intramembranous bone collar (**Figure 5C-E**). Maternal exercise did not significantly alter medullary vessel area or density (**Figure 5F-H**), but significantly increased bone collar vessel area and density (**Figure 5I-J**). Maternal exercise also significantly increased vessel area and density in the metaphyseal region, which captures the looping vessels along the chondro-osseous junction (**Figure 5K-L**). Together, these data suggest that maternal exercise enhanced growth plate remodeling and neovascularization, but did not directly stimulate osteogenesis, per se.

**Figure 5.**
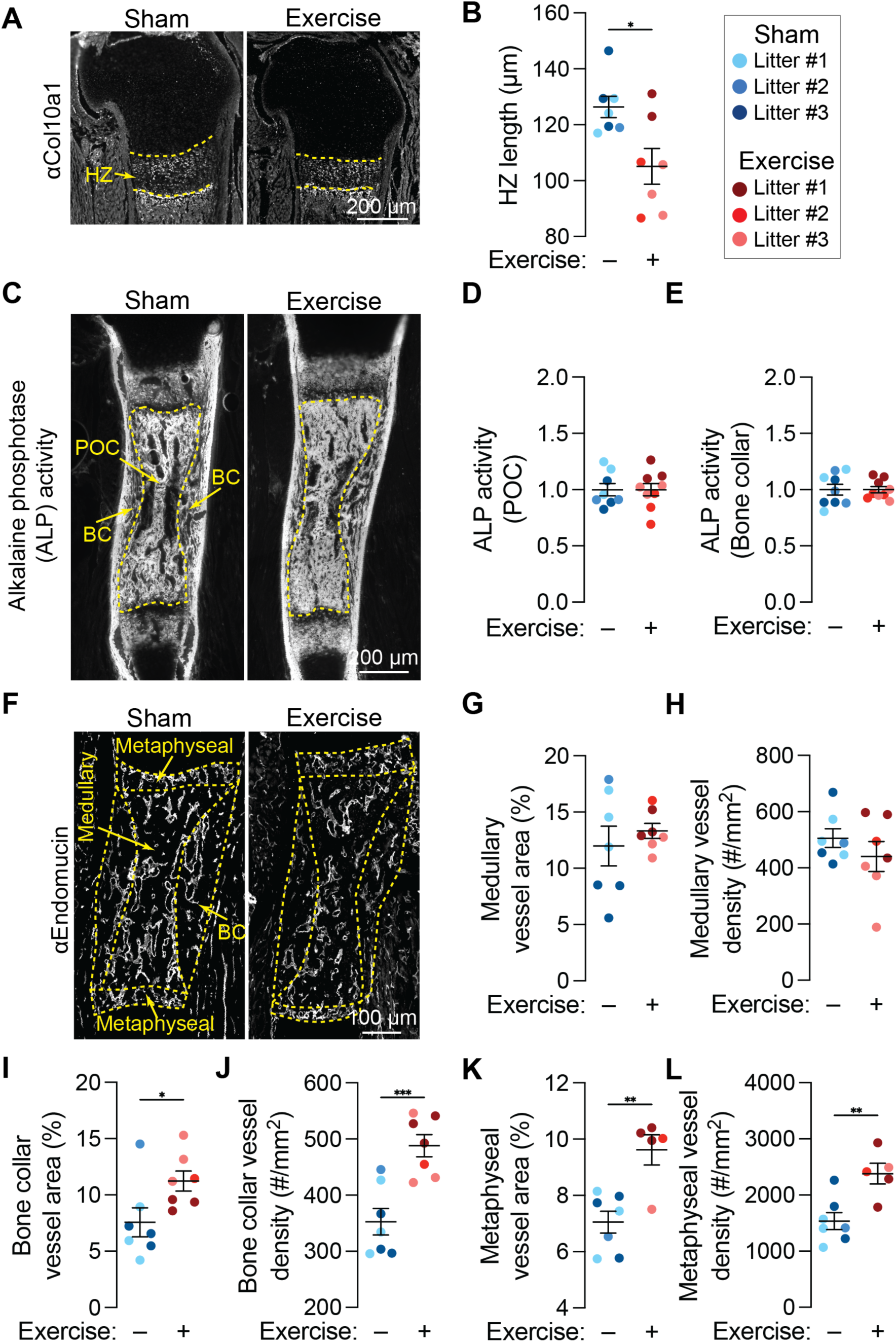
Effects of maternal exercise on growth plate morphometry, alkaline phosphatase (ALP) activity, and neovascular morphometry. **(A)** Fluorescent staining for collagen type 10 alpha 1 chain (Col10a1) in the proximal hypertrophic zone (HZ) of embryonic day 17.5 (E17.5) humeri. Scale bar = 200 µm. **(B)** HZ length quantification. **(C)** Fluorescent staining for ALP activity in E17.5 humeri. Scale bar = 200 µm. **(D)** Bone collar (BC) ALP activity and **(E)** primary ossification center (POC) ALP activity quantifications. **(F)** Fluorescent staining for blood vessels (endomucin) in E17.5 humeri. Scale bar = 100 µm. **(G)** BC vessel area, **(H)** BC vessel density, **(I)** medullary vessel area, **(J)** medullary vessel density, **(K)** metaphyseal vessel area, and **(L)** metaphyseal vessel density quantifications. Fetuses from the same litter are marked by the same color datapoint. * = *p*<0.05 using a Student’s t-test.

### 2.4 Effects of maternal exercise on fetal akinesia-impaired limb development

We next tested whether maternal exercise would influence joint and bone formation in the Splotch-delayed (Sp^d^) model of fetal akinesia.^3^ The Sp^d^ mouse exhibits limited muscle development and lacks spontaneous fetal movements;^32–34^ thus, this model has intact maternofetal transport, but altered mechanical loading. In these experiments, female dams pregnant with wild-type and Sp^d^ embryos were exercised from E13.5 to E15.5, inclusive, with harvest at E16.5. Like previous experiments, dams in the Sham group had access to locked wheels to prevent running.

Maternal exercise stimulated elbow joint morphogenesis in both wild-type and Sp^d^ fetuses, rescuing fetal akinesia-impaired joint parameters (**Figure 6A-B**). Specifically, fetal akinesia significantly reduced humeral medial condyle width and height (**Figure 6C-D**) and lateral condyle width and height (**Figure 6F-G**), but these defects were rescued in Sp^d^ embryos exposed to maternal exercise. Medial and lateral condyle depth were not significantly reduced by fetal akinesia, but were significantly increased by maternal exercise in Sp^d^ fetuses (**Figure 6E&H**).

**Figure 6.**
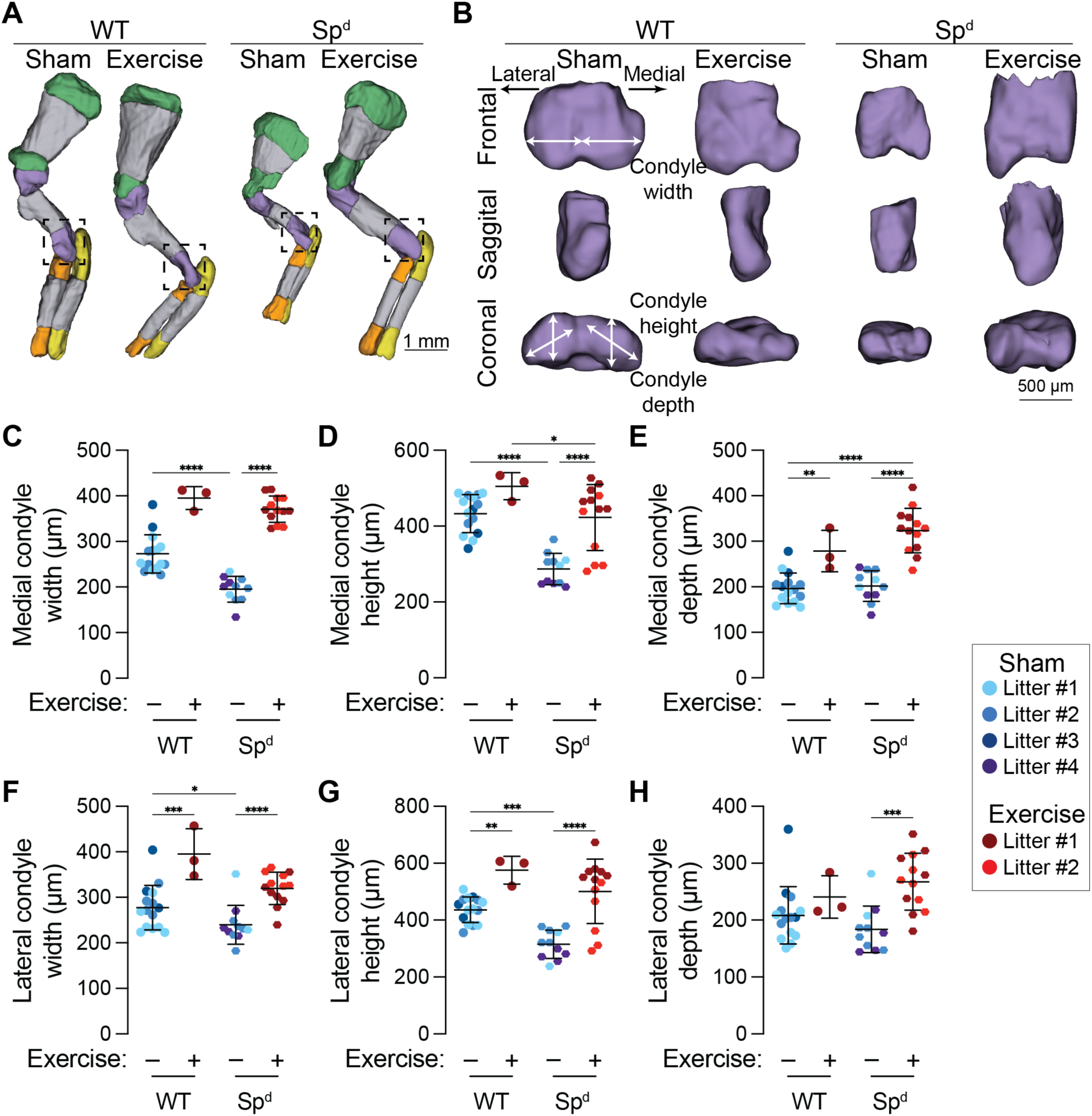
Effects of maternal exercise on fetal akinesia-impaired joint morphogenesis. **(A)** Optical projection tomography (OPT) reconstructions of embryonic day 16.5 (E16.5) forelimbs in wild-type (WT) and Splotch-delayed (Sp^d^) mice. Cartilaginous regions of the scapula, humerus, radius, and ulna are marked in green, purple, orange, and yellow, respectively. Ossified regions are marked in gray. **(B)** Zoomed projections of the cartilaginous distal humerus. **(C)** Medial condyle width, **(D)** medial condyle height, **(E)** medial condyle depth, **(F)** lateral condyle width, **(G)** lateral condyle height, and **(H)** lateral condyle depth quantifications. Fetuses from the same litter are marked by the same color datapoint. * = *p*<0.05 using a two-way ANOVA with Bonferroni correction.

Fetal akinesia significantly reduced humerus rudiment length, mineralized length, and mineralization ratio, but these defects were also rescued by maternal exercise (**Figure 7A-D**). Similar to joint morphogenesis, maternal exercise rescued these key bone morphogenesis outcomes. Lastly, we measured the effects of fetal akinesia and maternal exercise on morphogenesis of the deltoid tuberosity. Previous studies showed that the cartilaginous rudiment of the deltoid tuberosity forms in Sp^d^ mice at E14.5, but resorbs by E18.5 due to the absence of stimulatory mechanical cues.^44^ Consistently, we observed significantly reduced deltoid tuberosity volume in Sp^d^ fetuses at E16.5 (**Figure 7E**). However, maternal exercise rescued deltoid tuberosity volume to the level of wild-type fetuses. Importantly, the maternal exercise-induced rescue of deltoid tuberosity morphogenesis was preserved after controlling for overall rudiment growth (*i.e.,* after normalizing by rudiment length) (**Figure 7F**).

**Figure 7.**
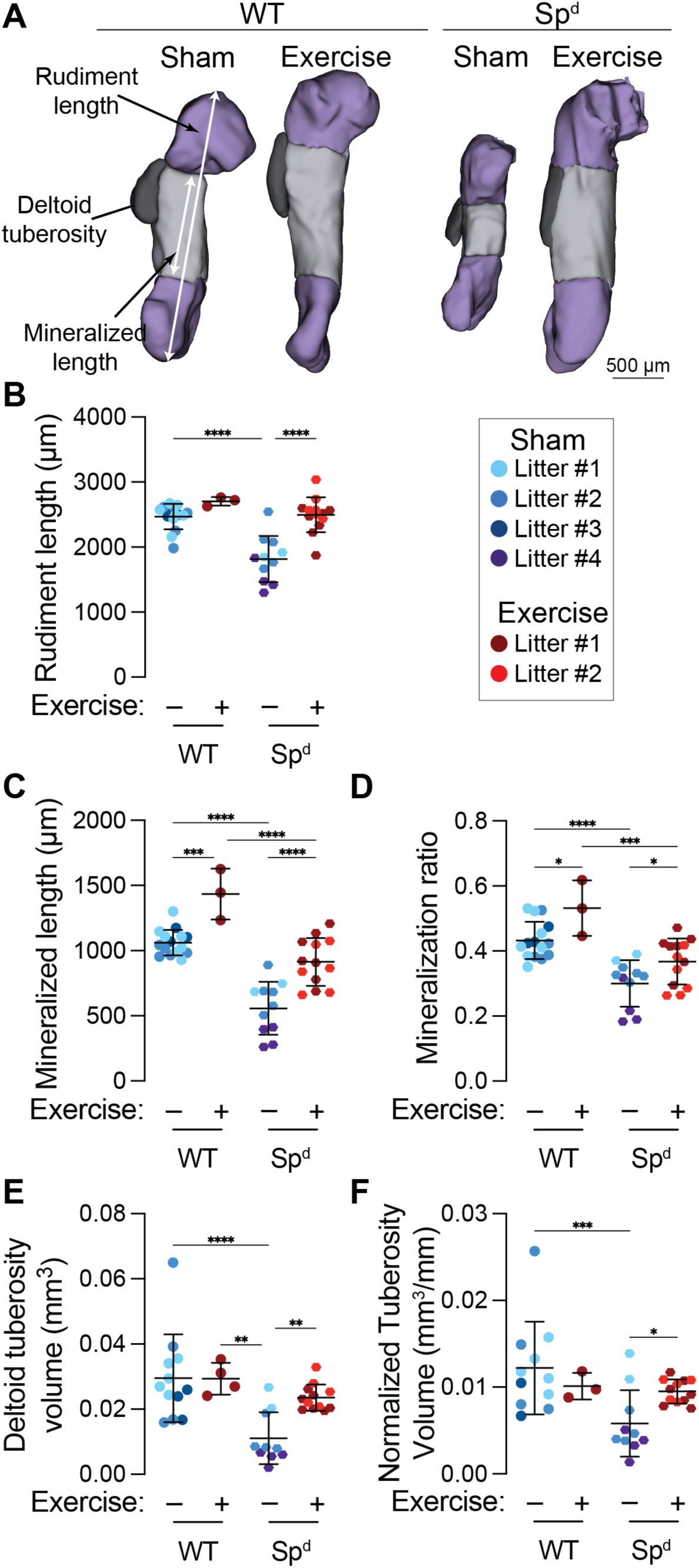
Effects of maternal exercise on fetal akinesia-impaired bone morphogenesis. **(A)** Optical projection tomography (OPT) reconstructions of embryonic day 16.5 (E16.5) humeri in wild-type (WT) and Splotch-delayed (Sp^d^) mice. The rudiment length and mineralized length are marked with white arrows. The deltoid tuberosity is contoured in black. Scale bar = 500 µm. **(B)** Rudiment length, **(C)** mineralized length, **(D)** mineralization ratio, **(E)** deltoid tuberosity volume, and **(F)** rudiment length-normalized deltoid tuberosity volume quantifications. Fetuses from the same litter are marked by the same color datapoint. * = *p*<0.05 using a two-way ANOVA with Bonferroni correction.

Finally, we tested the effects of direct mechanical loading on joint morphogenesis, in the absence of systemic maternal factors, by using *ex utero* mechanical stimulation bioreactor culture. Briefly, E15.5 embryonic forelimbs from wild-type and Sp^d^ fetuses were explanted and cultured for 7 days in a dynamic loading bioreactor. ^49,50^ Contralateral static control limbs from each genotype were cultured in the bioreactor without cyclic loading (**Figure 8A**). Under static conditions, wild-type and Sp^d^ joints were not significantly different in condyle height or condyle area (**Figure 8B-D**). This suggests that unloaded WT limbs exhibited the same impaired joint morphogenesis as Sp^d^ limbs due to explantation-induced arrest of active movement. However, dynamic loading significantly increased condyle height and condyle area, in both wild-type and Sp^d^ limbs.

**Figure 8.**
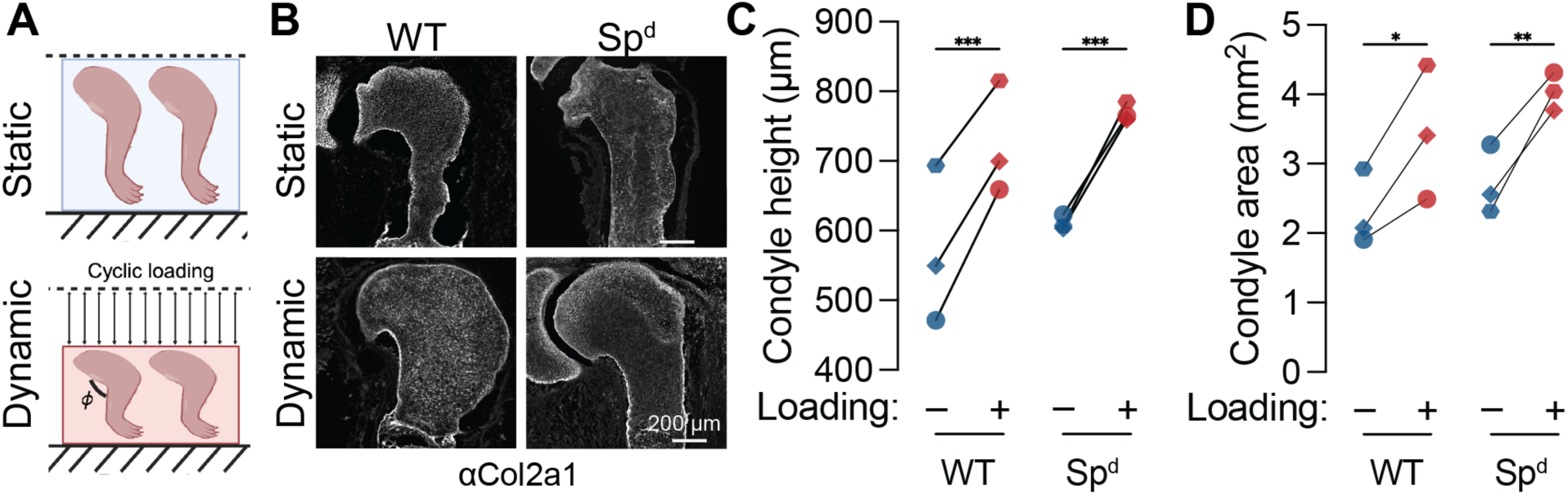
Effects of ex utero mechanostimulation bioreactor culture on fetal akinesia-impaired joint morphogenesis. **(A)** Schematic representation of ex utero mechanostimulation bioreactor culture of embryonic forelimbs from wild-type (WT) and Splotch-delayed (Sp^d^) mice. **(B)** Fluorescent staining for collagen type 2 alpha 1 chain (Col2a1) in the proximal joint of ex utero forelimbs. Scale bar = 200 µm. **(C)** Condyle height and **(D)** condyle area quantifications. Limbs from the same fetus are marked by the same shape and connecting line. * = *p*<0.05 using a two-way ANOVA with Bonferroni correction.

## 3.0 Discussion

Here, we describe the effects of maternal wheel running exercise on fetal skeletal development in mice. In wild-type fetuses, maternal exercise stimulated joint morphogenesis, bone growth, deltoid tuberosity formation, chondro-osseous junction remodeling, and blood vessel invasion. While maternal exercise increased the FW:PW ratio, an approximate marker of placental transport efficiency, we did not find other placental indicators of increased transport. Meanwhile, maternal exercise increased fetal bone expression of the YAP mechanotransducer target gene, Cyr61, and rescued joint morphogenesis, bone growth, and deltoid tuberosity formation in the Sp^d^ mouse model of fetal akinesia. Direct application of passive limb movement in explanted limbs, removed from all systemic factors, similarly stimulated joint morphogenesis. Taken together, our data suggest that the effects of maternal exercise on the developing skeleton are mediated, at least in part, by passive mechanical stimulation, but do not rule out contributions of maternofetal communication. These findings demonstrate that maternal exercise regulates fetal skeletal development, implicate maternal exercise as a platform for studying developmental mechanobiology of the skeleton *in vivo*, and motivate continued study into maternal exercise as a potential *in utero* intervention for fetal hypokinesia.

Maternal exercise can benefit offspring development. Controlled trials in humans showed that moderate maternal exercise (40-60% of maximum oxygen consumption (VO_2_ max), up to 50 min/day, 3 days/week) significantly increased fetal cardiac function at 36 weeks gestation^75^ and increased offspring brain maturity at 15 days old.^76^ Corroborating enhanced cognitive function outcomes, a preclinical rat study found that moderate-intensity maternal treadmill exercise (up to 8 m/min, 30 min/day, 5 days/week) significantly increased spatial learning memory in offspring.^77^ Preclinical models have also demonstrated the benefits of maternal exercise on offspring liver function. Specifically, moderate-intensity maternal treadmill exercise (up to 21 m/min, up to 60 min/day, 6 days/week) protected rat offspring from liver dysfunction associated with maternal high-fat/high-sugar diet^78^ and voluntary maternal wheel running exercise protected mouse offspring from developing non-alcoholic fatty liver disease associated with maternal high-fat diet.^79^ Voluntary maternal wheel running exercise in mice also protect offspring from developing mammary tumors after carcinogen challenge.^80^ Some studies have also described that maternal exercise prior to conception may benefit pregnancy and offspring.^81^ In our study, all mice, including sham and exercise groups, received the same *ad libitum* wheel acclimation prior to pregnancy. We used a targeted exercise approach, in which we exposed pregnant dams to one hour of supervised wheel running each day (broken in four 15-min bouts), for up to four days during the key window of fetal joint and bone development. We selected this limited regimen to mimic an achievable exercise program and because we observed that pregnant dams progressively diminish their running distances as pregnancy progresses when provided wheels *ad libitum*. Our findings contribute to a growing body of maternal exercise literature and pose new questions about how maternal exercise affects orthopaedic tissues.

We do not yet understand how maternal exercise affects offspring skeletal development. To date, only three preclinical studies have measured effects of maternal exercise on offspring skeletal development, but all focused on postnatal skeletal outcomes. One study found that voluntary wheel running exercise by pregnant mice, throughout gestation, increased osteogenic gene expression in the offspring femora, measured at two-months old.^82^ Conversely, another study found that subjecting pregnant rats to regular squat exercise throughout pregnancy (*i.e.,* food and water were raised so pregnant rats needed to stand on fully extended hindlimbs to eat and drink) significantly reduced tibial bone mineral density in the offspring, measured at four months old.^83^ The final study found that mild maternal treadmill exercise (30% VO_2_ max, unspecified exercise frequency) did not significantly alter femur length or bone mineral content in rat offspring at three-months old.^84^ In synthesis, these findings suggest that intensity and frequency of maternal exercise can regulate offspring skeletal health: too little exercise may not provide adequate stimuli, but too much may be detrimental. A recent systematic review and meta-analysis reported no significant difference in birthweight of babies born from mothers who completed vigorous intensity exercise in the third trimester compared with controls, but observed a small but significant increase in gestational age at delivery and a decrease in risk of prematurity.^85^ Our findings, using preclinical mouse models, suggest that moderate exercise during late gestation stimulates fetal skeletal development, but further studies are required to assay long-term benefits.

Synthesizing our findings, we posit a direct, if not sufficient, role of passive mechanical stimulation in joint and bone morphogenesis. This is based on several lines of experimental reasoning. First, we observed effects of maternal exercise on YAP/TAZ-target gene expression, growth plate remodeling, and vascular morphogenesis at the chondro-osseous junction, which are consistent with our prior findings that YAP and TAZ mediate mechanical control of limb development and regulate neovascular invasion-mediated growth plate remodeling.^53^ Further work will be needed to test for direct roles of YAP/TAZ mechanotransduction in maternal exercise-induced limb morphogenesis. Second, maternal exercise substantially rescued joint and bone morphogenesis in the Sp^d^ model of fetal akinesia, which disrupts fetal movement without altering maternofetal communication. Notably, maternal exercise prevented akinesia-induced resorption of the deltoid tuberosity, an established hallmark of mechanoregulated morphogenesis.^35,57^ Normalizing deltoid tuberosity volume by rudiment length, to control for overall rudiment growth induced by maternal exercise, did not abrogate the rescue, further suggesting a local mechanical response. Lastly, we used an orthogonal *ex utero* approach for direct mechanical stimulation of explanted limbs, removed from systemic factors.^49–51,53,54^ These data show that mechanical cues are sufficient to stimulate joint morphogenesis in both wild-type and Sp^d^ mice.

Our data do not preclude the contribution of maternal-derived humoral signals. For example, a rat study found that moderate maternal treadmill exercise (up to 17 m/min, up to 50 min/day, 7 days/week) increased fetal weight and placental transport efficiency via increased IGF signaling.^67^ The role of IGF signaling in maternal exercise is plausible since dysregulation of the IGF system has been implicated in the pathogenesis of fetal growth restriction in mice,^86–88^ rats,^89,90^ rabbits,^91^ sheep,^92–94^ and humans.^95,96^ We observed increased FW:PW ratio, which is indicative of increased placental transport, but did not observe changes in placental zone morphometry,^70,71^ placental IGF-1 levels,^72,73^ or nutrient transporter expression.^70,71^ This may stem from different intensity (*i.e.,* wheel running vs. treadmill) and duration (*i.e.,* 3-4 days vs. 20 days) in our exercise modality. Although we do not observe evidence of increased IGF signaling in our exercise model, it is possible that other maternally secreted humoral signals are involved. For example, a mouse study found that moderate maternal treadmill exercise (up to 40-65% VO_2_, up to 60 min/day, 7 days/week) enhanced fetal muscle development via elevated apelin signaling, which demethylated the promoter region of *Ppargc1a* and promoted mitochondriogenesis.^68^ Future studies will be required to determine these putative contributions.

This study has limitations. First, our study does not vary exercise duration or intensity. Human pregnancy guidelines recommend that maternal exercise can be beneficial through means that depend on the intensity, duration, and exercise modality.^62^ Although the mice in our study ran consistent distances, our supervised wheel running approach could not modulate exercise intensity. Future studies using controlled treadmill running and longitudinal endpoint assessments of maternal exercise physiology will be required. Second, we utilize the Sp^d^ mouse model of fetal akinesia, which produces fetuses that have no mature skeletal muscle. Though this mouse represents an extreme case of absent fetal movements, the Sp^d^ model prevented us from studying possible influences of trophic factors from fetal muscle in the response to maternal exercise. In future studies, we aim to use the *mdg* mouse model of fetal akinesia as a more clinically relevant model of amyoplasia that may preserve bone-muscle crosstalk. Lastly, though our evidence to suggests that maternal exercise regulates fetal skeletal development in part through passive restoration of mechanical signals, this study was not designed to define the underlying mechanotransductive mechanisms and our data do not eliminate the possible contributions of altered maternofetal communication. Likewise, we do not yet know whether the developing cell populations affected by fetal akinesia are directly stimulated by maternal exercise, or whether the observed rescue effects are mediated by indirect cellular mechanisms.

Taken together, these findings identify maternal exercise as a regulator of fetal skeletal development, providing a platform for studying skeletal developmental mechanobiology and suggesting potential therapeutic applications for fetuses with impaired movement.

## 4.0 Materials and methods

### 4.1 Animal husbandry and care

C57BL/6J male and female mice were purchased from the Jackson Laboratory (Jackson Laboratory, Bar Harbor, ME, USA). Mice were housed and bred for experiments at the University of Pennsylvania. Heterozygous Splotch-delayed (Pax3^spd/+^) male and female mice were also purchased from the Jackson Laboratory. These mice were housed and bred at Imperial College London. Sp^d^ embryos were genotyped using PCR on DNA derived from embryonic head tissue. DNA underwent 30 30 second reaction cycles at 94°C, 60°C, and 74°C. For controls, the following primers were used: 5’ AGGGCCGAGTCAACCAGCACG 3’ and 3’ CACGCGAAGCTGGCGAGAAATG 5’. For mutants, the following primers were used: 5’ AGTGTCCACCCCTCTTGGCCTCGGCCGAGTCAACCAGGTCC 3’ and 3’ CACGCGAAGCTGGCGAGAAATG 5’

Prior to maternal exercise experiments, all mice were fed regular chow (Purina LabDiet, St. Louis, MO, USA) *ad libitum* and housed in cages containing 2–4 animals each. Mice were maintained at constant 25°C on a 12 hr light/dark cycle. Protocols were approved by the Institutional Animal Care and Use Committees at the University of Pennsylvania for experiments conducted in the United States (Protocol no: 806482) and United Kingdom experiments were conducted under the UK Home Office Project License number PPL P39D18B9C. All US-based animal procedures were performed in adherence to United States federal guidelines of animal care. All UK-based procedures were performed in accordance with the UK Animals Scientific Procedures Act of 1986 and were approved by the institutional Animal Welfare and Ethical Review Body (AWERB). Additionally, experiments were conducted in accordance with the 3R principles in animal research.^97–99^

### 4.2 Maternal exercise

Female C57BL/6J mice were housed individually with the Mouse Home Cage Running Wheel system (Columbus Instruments, Columbus, OH) for at least two weeks prior to timed matings. This free-spinning wheel running system allowed us to measure voluntary running activity during the acclimation period. After timed pregnancies, mice were individually housed, without running wheels, from E0 to E13.5. Mated females were weighed at E0 and E12.5 to confirm pregnancy by weight gain. For *ad libitum* wheel running experiments, dams pregnant with E13.5 embryos were re-housed with their running wheels and allowed to run freely from E13.5 through E17.5, with harvest at E17.5. For supervised wheel running experiments, mice were individually housed without running wheels from E13.5 through E17.5, but exercised daily from E13.5 to E16.5, inclusive (**Figure S1A**). Each day, the pregnant dam was transferred back to her cage with the running wheel and placed onto the wheel. To prevent encourage running, the wheel was lifted slightly off the ground for each bout of exercise. Mice exercised for 15 min, then rested for 15 min with the wheel back in the cage. This process was repeated four times, so that dams received 1 hr of encouraged wheel running exercise, per day. At the end of this exercise regimen, pregnant dams were returned to their cages without wheels. Sham mice were included in *ad libitum* and supervised wheel running experiments, which received the same access to wheels, but the wheels were locked to prevent running.

Humane endpoints were predefined, including criteria such as poor grooming, sunken eyes, hunched posture, severe axial deviation, lack of food and water intake, significant weight loss, bloody feces, severe respiratory issues, debilitating diarrhea, seizures, paresis, and abscesses. No adverse effects of exercise were observed in any of the mice and humane endpoints were reached during the study. Pregnant dams were monitored closely until euthanasia, when embryos reached E17.5.

*Ad libitum* wheel access during the maternal exercise period yielded highly variable running distances and low running distances at later timepoints that are important for joint and bone development (**Figure S1B-D**). Thus, Sp^d^ mice were only exercised using supervised wheel running. Sp^d^ mice underwent the same procedures described above, except embryos were harvested at E16.5.

### 4.3 Microcomputed tomography (µCT)

The Microct45 system (SCANCO Medical Ag, Brüttisellen, Switzerland) was used to capture high-resolution three-dimensional (3D) microcomputed tomography (MicroCT) images of fixed E17.5 forelimbs from C57BL/6J embryos. E17.5 forelimbs were submerged in phosphate buffered saline (PBS, Thermo Fisher Scientific, Waltham, MA, USA) for image capture. The following imaging parameters were applied: isotropic voxel size of 3 µm, integration time of 300 ms, x-ray intensity of 114 µA, and peak tube voltage of 70 kVp. A three-dimensional Gaussian filter of 1.2 with filter support of 2 was used for noise suppression, and mineralized tissue was segmented from the PBS or soft tissue using a threshold of 140 mgHA/cm^3^. The acquired images were analyzed using the manufacturer-provided software.

### 4.4 Optical projection tomography (OPT)

Right forelimbs from E17.5 C57BL/6J embryos and E16.5 Sp^d^ embryos were harvested for skeletal preparations with Alcian blue (MilliporeSigma, Billerica, MA, USA) and Alizarin red (MilliporeSigma).^100^ Harvested forelimbs were frozen at −20°C prior to preparation. Upon thawing, samples were dissected to remove the outer epidermal layer, which improves stain penetration, then incubated in 95% (v/v) ethanol (Decon Laboratories Inc., Swedeland, PA, USA) overnight. The next day, samples were washed with acetone (MilliporeSigma) for 1-2 hr, then incubated with acetone overnight. After acetone incubation, samples were rinsed with deionized water (ddH_2_O) for 1-2 hr, then incubated for 24 hr in a 150 mg/L Alcian blue solution (80% ethanol (v/v), 20% acetic acid (v/v) (Thermo Fisher Scientific)). The following day, samples were washed with 70% ethanol (v/v) four times for 1 hr each, washed for 1 hr with 1% potassium hydroxide (KOH, w/v) (VWR International, Radnor, PA, USA), then incubated overnight in a 50 mg/L Alizarin red solution (1% KOH (w/v), ddH_2_O). After Alizarin red incubation, the stain was gently poured off and samples were placed in a storage mixture (50% ethanol (v/v), 50% glycerol (v/v) (MilliporeSigma) and stored at 4°C until embedding for optical projection tomography (OPT). All washes and incubations prior to the KOH wash were carried out at room temperature (RT) with rocking. Since KOH causes the skeletal preparations to be fragile, the KOH wash and Alizarin red incubation were conducted at RT without rocking.

Forelimbs stained with Alizarin red and Alcian blue were embedded in 1% agarose (w/v) (MilliporeSigma) hydrogels for optical projection tomography (OPT) imaging (**Figure S2**).^101,102^ To prepare the 1% agarose hydrogels, agarose powder was dissolved into boiling ddH_2_O, then sterile filtered using the Steriflip® (MilliporeSigma). The sterilized 1% agarose solution was poured into 35 mm petri dishes and samples were carefully placed into the polymer solutions at medium depth. Embedded samples were allowed to solidify on ice. Excess agarose was removed for all samples, then samples were dehydrated for 24 hr in methanol (Thermo Fisher Scientific). To ensure full dehydration, the methanol solution was replaced 2-3 times during the 24 hr incubation. The following day, samples were cleared for 24 hr using a solution of 50% benzoic acid (v/v) (MilliporeSigma) and 50% benzyl benzoate (v/v) (MilliporeSigma). For OPT imaging, samples were staged using a custom OPT imaging apparatus,^103^ which captures 400 images of each forelimb as it is rotated 360°. Image sets were reconstructed into two-dimensional (2D) sections using NRecon software (Micro Photonics Inc., Allentown, PA, USA), cropped using ImageJ/Fiji software (National Institutes of Health, Bethesda, MD, USA), then segmented using ITK SNAP (Penn Image Computing and Science Laboratory, Philadelphia, PA USA). Measurements of rudiment lengths, mineral lengths, and joint shape features were quantified from 3D reconstructions using Blender (Blender Foundation, Amsterdam, NL) and 3D Slicer.^104^

### 4.5 Cryohistology and immunofluorescence staining

C57BL/6J embryos harvested at E17.5 were fixed in 4% paraformaldehyde (PFA, Electron Microscopy Sciences, Hartfield, PA, USA) at 4°C overnight, then transferred to 30% sucrose (w/v) (Thermo Fisher Scientific) in PBS for 24 hr. After sucrose infiltration, embryonic forelimbs were isolated and embedded in optical cutting temperature (OCT) compound (Tissue-Tek, Torrance, CA, USA) for cryosectioning with an Epredia^TM^ CryoStar^TM^ NX70 Cryostat (Thermo Fisher Scientific). 10 µm thick tissue sections were obtained on cryotape (Section Lab Co, Hiroshima, Japan), as described in detail in a previously published protocol.^105^ Tape sections were glued to microscope slides with Norland Optical Adhesive 81 (Norland Products Inc., Jamesburg, NJ, USA), then processed using standard kits and immunofluorescent protocols. For all protocols, samples were mounted using Fluoromount-GT (Thermo Fisher Scientific), imaged using the ZEISS AxioScan.Z1 Slide Scanner (ZEISS, Oberkochen, Germany), and analyzed using ImageJ/Fiji.

Alkaline phosphatase (ALP) staining was conducted using the Vector blue Alkaline Phosphatase (ALP) substrate kit (SK-5300, Vector Laboratories, Newark, CA, USA) according to the manufacturer’s instructions.

Phalloidin staining with Hoechst counterstain (Thermo Fisher Scientific) was conducted using Alexa Fluor 647 Phalloidin (A22287, Thermo Fisher Scientific) according to manufacturer’s instructions.

Immunofluorescent imaging of Collagen 10, Endomucin, Yes-associated protein (YAP), and Connective tissue growth factor-like protein 1 (Cyr61/CCN1) was conducted using standard immunofluorescent protocols. Briefly, glued tissue sections were rehydrated in PBS, then blocked with 5% goat serum (MilliporeSigma) in 0.3% Triton-X-100 (MilliporeSigma) for 30 min. The following primary antibodies were applied overnight at 4°C (dilutions and catalog numbers provided): rabbit polyclonal anti-Collagen 10 (ab182563, Abcam, Cambridge, UK), rat monoclonal anti-Endomucin (1:100, sc-65495, Santa Cruz Biotechnology, Dallas, TX, USA), rabbit monoclonal anti-YAP (1:100, 14074, Cell Signaling Technologies, Danvers, MA, USA), rat monoclonal anti-Cyr61 (1:20, MAB4864, R&D Systems, Minneapolis, MN, USA). After primary antibody incubation, samples were washed thrice with PBS, then incubated with the following secondary antibodies for 2 hr at RT: goat anti-rat 647 (1:1000, A-21247, Thermo Fisher Scientific), goat anti-rat 488 (1:1000, A-11006, Thermo Fisher Scientific), goat anti-rabbit A647 (1:1000, A-27040, Thermo Fisher Scientific), and goat anti-rabbit A488 (1:1000, A11034, Thermo Fisher Scientific). Hoechst counterstain (1:2000, Thermo Fisher Scientific), was added during the secondary antibody incubation. Samples were then washed thrice with PBS prior to mounting.

### 2.6 Paraffin histology and tinctorial staining

Placentas were isolated during embryo harvests, flash frozen in liquid nitrogen, and stored at - 80°C until paraffin histology. Half of each placenta was thawed and processed for paraffin histology using a Thermo Scientific Excelsior AS Tissue Processor (Thermo Fisher Scientific). Processed samples were paraffin-embedded and section to 10 µm using a Leica RM 2030 Microtome (Leica, Bala Cynwyd, PA, USA). Sections were stained for hematoxylin and eosin using standard protocols. Stained sections were imaged using a ZEISS Axio Scan.Z1 Slide Scanner and analyzed using ImageJ/Fiji.

### 4.6 Protein lysis, enzyme-linked immunosorbent assay (ELISA), and western blot

The remaining half of each placenta was processed for enzyme-linked immunosorbent assay (ELISA) and western blot. Approximately 20 mg of placental tissue was placed in 550 µL of 1X Cell Lysis Buffer (R&D Systems) and homogenized for 5 min at max speed using the Bullet Blender Storm Tissue Homogenizer (Next Advance, Troy, NY, USA). Homogenized lysates were incubated on ice for 15 min, transferred to a clean 1.5 mL centrifuge tube, and centrifuged at 13,000 g for 10 min at 4°C using a Sorvall Legend Micro 17R centrifuge (Thermo Fisher Scientific). Protein levels were quantified using a Pierce^TM^ BCA Protein Assay Kit (23227, Thermo Fisher Scientific) according to manufacturer’s instructions.

Placental levels of Insulin-like growth factor-1 (IGF-1) were quantified using the Mouse IGF-1 ELISA Kit (RAB0229, Thermo Fisher Scientific) according to manufacturer’s instructions.

Placental levels of Glucose transporter 1 (GLUT1), Sodium-coupled neutral amino acid transporter 4 (SNAT4), and Fatty acid transporter 4 (FATP4) were quantified using standard western blot protocols. Briefly, protein solutions were diluted to 20 µg of protein in 20 µL of RNAse-free H_2_O (Thermo Fisher Scientific), then mixed in a 1:1 ratio with a solution of 95% 2X

Laemmli Buffer (v/v) (Bio-Rad, Philadelphia, PA, USA) and 5% *β*-mercaptoethanol (v/v) (Bio-Rad). Samples were loaded onto 4-15% Mini-PROTEAN® TGX^TM^ Precast Protein Gels (Bio-Rad) with the Precision Plus Kaleidoscope ladder (Bio-Rad) and run using the Mini-PROTEAN® Tetra System (Bio-Rad). Bands were transferred to Immun-Blot® PVDF Membranes (Bio-Rad) using the Trans Blot® Turbo^TM^ Transfer System (Bio-Rad). Membranes were washed thrice in tris-buffered saline with Tween 20 (TBST, Bio-Rad), then blocked using Blotting-Grade blocker (Bio-Rad) 1.5 hr on a rocker at RT. The following primary antibodies were applied overnight at 4°C with rocking (dilutions and catalog numbers provided): rabbit monoclonal anti-GLUT1 (1:1000, 12939, Cell Signaling Technologies), rabbit polyclonal anti-SLC38A4/SNAT4 (1:1000, 20857-1-AP, Proteintech, Rosemont, IL, USA), rabbit monoclonal anti-SLC27A4/FATP4 (1:1000, ab200353, Abcam), rabbit polyclonal anti-*β*-actin (1:1000, Cell Signaling Technologies), rabbit polyclonal anti-histone H3 (1:5000, Thermo Fisher Scientific). After primary antibody incubation, samples were washed thrice with TBST, then incubated with anti-rabbit IgG, HRP-linked antibody (1:1000, 7074P2, Cell Signaling Technologies) for 1.5 hr at RT on a rocker. Bands were detected using the Amersham^TM^ ECL^TM^ Prime Western Blotting Detection Reagent (Thermo Fisher Scientific) according to manufacturers instructions, then imaged using the Bio-Rad ChemiDoc^TM^ MP Imaging System (Bio-Rad). Band intensities were quantified using ImageJ/Fiji.

### 4.7 Ex vivo bioreactor culture

Histological images generated by Ahmed et al.^54^ were reanalyzed using quantitative histomorphometry to determine the effects of *ex vivo* mechanical loading on embryonic limb development. Full materials and methods for generating these samples are provided in Ahmed et al. ^54^ Briefly, E15.5 forelimbs were harvested from WT and Sp^d^ embryos then cultured using the Ebers TC-3 bioreactor system (Don Whitley Scientific, Bingley, UK). Contralateral limbs were randomly assigned to either Static or Dynamic culture conditions. Static forelimbs were placed within the bioreactor foam supports without any loading. Dynamic forelimb explants were loaded at 0.67 Hz to a displacement of 2 mm for 2 h, 3 times per day. In previous studies, we demonstrated that this compression produces a cyclic flexion angle of approximately 14°, which led to the most physiological growth of tissues.^49^ All explanted limbs were cultured at air-liquid interface with osteogenic media (ɑ-MEM GlutaMAX supplemented with 1% penicillin-streptomycin with amphotericin B, 100 µM ascorbic acid, 2 mM *β*-glycerophosphate, and 100 nM dexamethasone) for 7 days at 37°C in normoxic conditions. Media was changed daily.

Cultured forelimbs were prepared for cryohistology. Full materials and methods for staining and imaging are provided in Ahmed et al.^10^ The following antibodies were used (dilutions and catalog numbers provided): mouse monoclonal anti-collagen type II (1:50, MAB8887, MilliporeSigma) and rabbit anti-mouse (Alexa Flur® 488) (1:200, ab150125, Abcam). Samples were imaged using either the ZEISS LSM 510 (ZEISS) or the Leica CF6 (Leica) confocal laser scanning microscope. Images were analyzed using ImageJ/Fiji.

### 4.8 Statistical analysis

The “pwr2” package in R (Indianapolis, IN, USA) was used to conduct power analysis for large effect sizes (⍺ = 0.05, **β** = 0.20, f = 0.4 –0.6). GraphPad Prism software Version 10.4.1 (GraphPad Software, San Diego, CA, USA) was used to conduct all statistical analyses. A Student’s t-test was used to determine significant differences for all outcome measurements in the C57BL/6J experiments. A two-way analysis of variance (ANOVA) with Bonferroni correction was used to determine significant differences for all outcome measurements in the *in vivo* Sp^d^ experiments. A repeated measures two-way ANOVA with Bonferroni correction was used to determine significant differences for all outcome measurements in the *ex utero* Sp^d^ experiments.

## 6.0 Acknowledgements

## 6.1 Funding

European Research Council (ERC) – European Union’s Seventh Framework Programme grant agreement number 336306 (NCN)

ERC – Horizon 2020 Research and Innovation Programme grant agreement number 101124603 (NCN)

Research Ireland grant 23/US/3923 (NCN)

NIH/NICHD grant R01-HD113596 (JDB, ND)

NIH/NIAMS grant P30-AR069619 (JDB, ND)

NIH/NIGMS grant K12GM081259 (CJP)

NSF CMMI grant 15-48571 (JDB)

University of Pennsylvania Center for Undergraduate Research and Fellowships (CURF) (CJP, DCG, ME, JDB)

### 6.2 Author contributions

Conceptualization: CJP, YH, NK, JDB, NCN

Methodology: CJP, YH, NK, DCG, ME, SA

Investigation: CJP, YH, NK, DCG, SA, ME

Visualization: CJP, YH, NK, DCG, ME, SA, JDB, NCN

Supervision: ND, RAS, JDB, NCN

Writing – original draft: CJP, JDB, NCN

Writing – review & editing: CJP, YH, NK, DCG, ME, SA, ND, RAS, JDB, NCN

## 7.0 Supplementary data

**Supplemental Figure 1.**
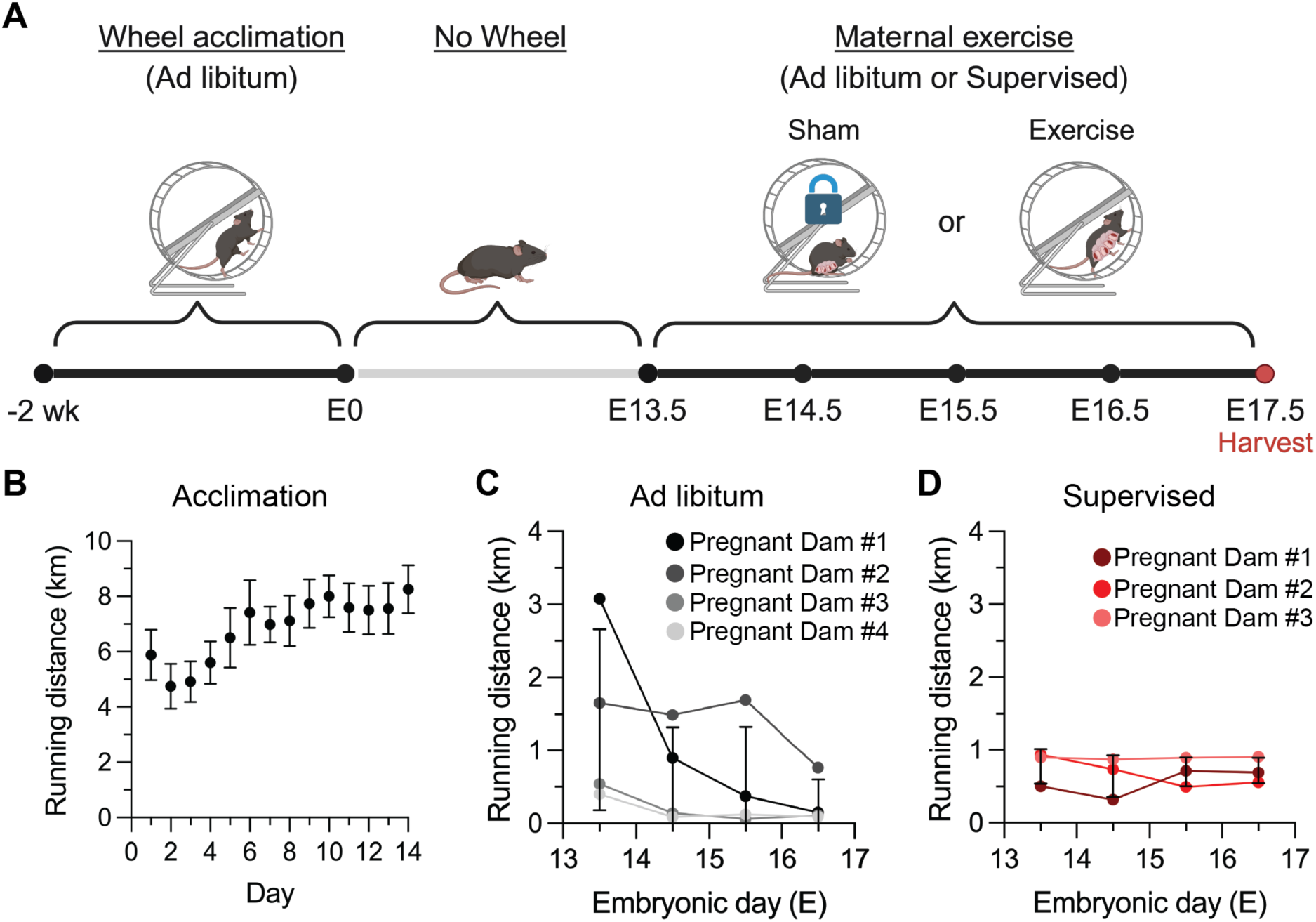
Wheel acclimation and maternal exercise regimen. **(A)** Schematic representation of maternal exercise modalities. Female mice were acclimated to running wheels with *ad libitum* access for two weeks prior to timed pregnancies. After timed pregnancies, pregnant dams had no wheel access until embryonic day 13.5 (E13.5). At E13.5, pregnant dams were assigned to one of two exercise modalities: (1) *Ad libitum* wheel access until harvest at E17.5 or (2) Supervised wheel running daily until harvest at E17.5. Each exercise modality included Sham mice, which had the same wheel exposure, but wheels were locked so that the pregnant dams could not use them to exercise. Embryos were harvested at E17.5 for downstream analyses. **(B)** Daily running distance throughout the two-week acclimation period. **(C)** Daily running distance for pregnant dams using the *Ad libitum* exercise modality. **(D)** Daily running distance for pregnant dams using the Supervised running exercise modality. Each line represents one pregnant dam.

**Supplemental Figure 2.**
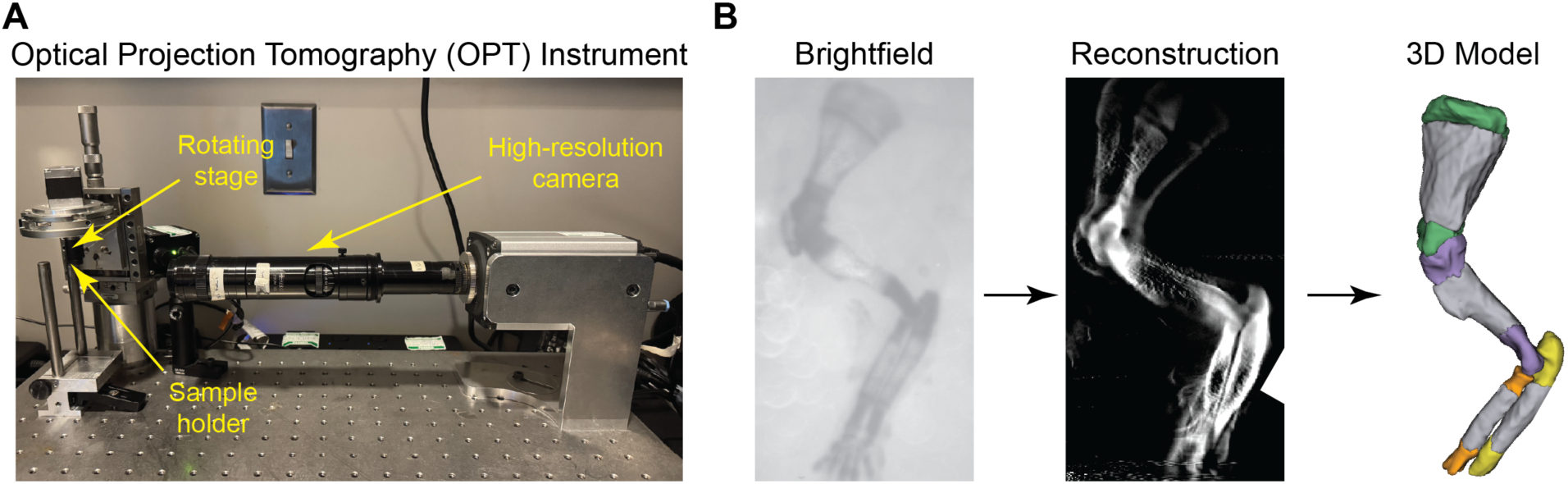
Optical projection tomography (OPT) imaging of fetal forelimbs. **(A)** OPT apparatus with high-solution camera, rotating stage, and magnetic sample holder marked by yellow arrows. **(B)** Example process for how a series of 400 brightfield images captured at 360° was reconstructed and converted into a 3D model.

